# Cerebral blood flow predicts differential neurotransmitter activity

**DOI:** 10.1101/207407

**Authors:** Juergen Dukart, Štefan Holiga, Christopher Chatham, Peter Hawkins, Anna Forsyth, Rebecca McMillan, Jim Myers, Anne R Lingford-Hughes, David J Nutt, Emilio Merlo-Pich, Celine Risterucci, Lauren Boak, Daniel Umbricht, Scott Schobel, Thomas Liu, Mitul A Mehta, Fernando O Zelaya, Steve C Williams, Gregory Brown, Martin Paulus, Garry D Honey, Suresh Muthukumaraswamy, Joerg Hipp, Alessandro Bertolino, Fabio Sambataro

**Affiliations:** F. Hoffmann-La Roche, pharma Research Early Development, Roche Innovation Centre Basel, Basel, Switzerland.; Department of Neuroimaging, Institute of Psychiatry, Psychology & Neuroscience, King’s College London, London, United Kingdom; School of Pharmacy, Faculty of Medical and Health Sciences, The University of Auckland, Auckland, New Zealand; Center for Functional MRI, University of California San Diego, 9500 Gilman Drive MC 0677, La Jolla, CA 92093, United States; Departments of Radiology, Psychiatry and Bioengineering, University of California San Diego, 9500 Gilman Drive, La Jolla, CA 92093, United States. Department of Psychiatry; University of California, San Diego, La Jolla, USA Veterans Affairs San Diego Healthcare System, San Diego, USA; Institute Of Psychiatry, Department of Basic Medical Science, Neuroscience and Sense Organs, University of Bari ‘Aldo Moro’; Department of Experimental and Clinical Medical Sciences (DISM), University of Udine, Udine, Italy; Neuropsychopharmacology Unit, Imperial College London, London, United Kingdom

## Abstract

Application of metabolic magnetic resonance imaging measures such as cerebral blood flow in translational medicine is limited by the unknown link of observed alterations to specific neurophysiological processes. In particular, the sensitivity of cerebral blood flow to activity changes in specific neurotransmitter systems remains unclear. We address this question by probing cerebral blood flow in healthy volunteers using seven established drugs with known dopaminergic, serotonergic, glutamatergic and GABAergic mechanisms of action. We use a novel framework aimed at disentangling the observed effects to contribution from underlying neurotransmitter systems. We find for all evaluated compounds a reliable spatial link of respective cerebral blood flow changes with underlying neurotransmitter receptor densities corresponding to their primary mechanisms of action. The strength of these associations with receptor density is mediated by respective drug affinities. These findings suggest that cerebral blood flow is a sensitive brain-wide in-vivo assay of metabolic demands across a variety of neurotransmitter systems in humans.

## 1. Introduction

Metabolic task-based and resting state magnetic resonance imaging (MRI) of blood oxygen level dependence and cerebral blood flow (CBF) are now commonly applied for studying human brain function, disease pathology and for evaluation of pharmacodynamic (PD) effects associated with pharmacological interventions ^1–9^. Both oxygen and glucose are delivered to brain structures by CBF to address their metabolic demands ^10^. In particular, state of the art arterial spin labeling (ASL) sequences now allow for quantitative CBF evaluation and are used for evaluation of metabolism associated with neural activity ^11–15^. Despite their wide-spread applications in healthy and diseased populations there is limited understanding of whether and how metabolic effects measured through these techniques reflect underlying activity in specific neurotransmitter systems^1,2,4,7,16^. A better understanding could unveil potential mechanisms of disease and crucial components of complex drug action that underpin functional modulation ^7,17^ Major limitations discussed in that context are potential contributions of confounding physiological, e.g. cardiovascular effects, unclear association with specific neurotransmitter systems and agonist and antagonist effects and a narrow cross-species translational value restricting comparisons to a descriptive anatomical level ^18–26^. Addressing those limitations is therefore key to wide-spread application of metabolic MRI in translational medicine.

Receptor theory provides a possible way of addressing these limitations ^7,27^ This theory posits that relationships between drug kinetics and observed PD effects depend on both the drug (i.e. receptor affinity and mechanism of action) and the biological system (i.e. receptor density and activity). Based on this concept drugs affecting specific receptor systems should lead to higher metabolic changes in regions with higher respective receptor densities. The strength of this relationship should be further dependent on the affinity of the compounds to the respective receptor systems. However, this assumption only partially holds for drugs with an indirect mechanism of action (i.e. allosteric modulators and reuptake inhibitors). These drugs do not directly induce activation but rather facilitate the effects of activation induced through other mechanisms. One would therefore expect their effects to co-localize with such underlying activity. In contrast, direct a(nta)gonists should additionally also activate yet inactive regions with high respective receptor densities.

Using this concept, we assess if ASL derived CBF changes (ΔCBF) induced by seven established compounds with known direct or indirect dopaminergic, serotoninergic, glutamatergic and/or GABAergic mechanisms of action (escitalopram, methylphenidate, haloperidol, olanzapine, low and highdose of risperidone, ketamine and midazolam, Table 1) are associated with respective receptor densities, underlying activity and affinities to the respective receptor types (Figure 1). Based on the above considerations, we hypothesize stronger ΔCBF in regions with higher respective receptor densities in particular for drugs with a direct mechanism of action and to a weaker extent for allosteric modulators and reuptake inhibitors. Furthermore, we expect stronger ΔCBF in regions with higher underlying activity for both compounds with direct (direct agonists and antagonists) and indirect mechanisms of action (allosteric modulators and reuptake inhibitors). Lastly, we hypothesize the association strength between ΔCBF and receptor densities to be dependent on the respective drug affinities. We test those hypotheses by first evaluating the link between drug-induced ΔCBF with ex vivo and in vivo estimates of different receptor densities and expected underlying activity. In a second step we then test if the association strength of these relationships is dependent on the respective drug affinities.

**Fig. 1.**
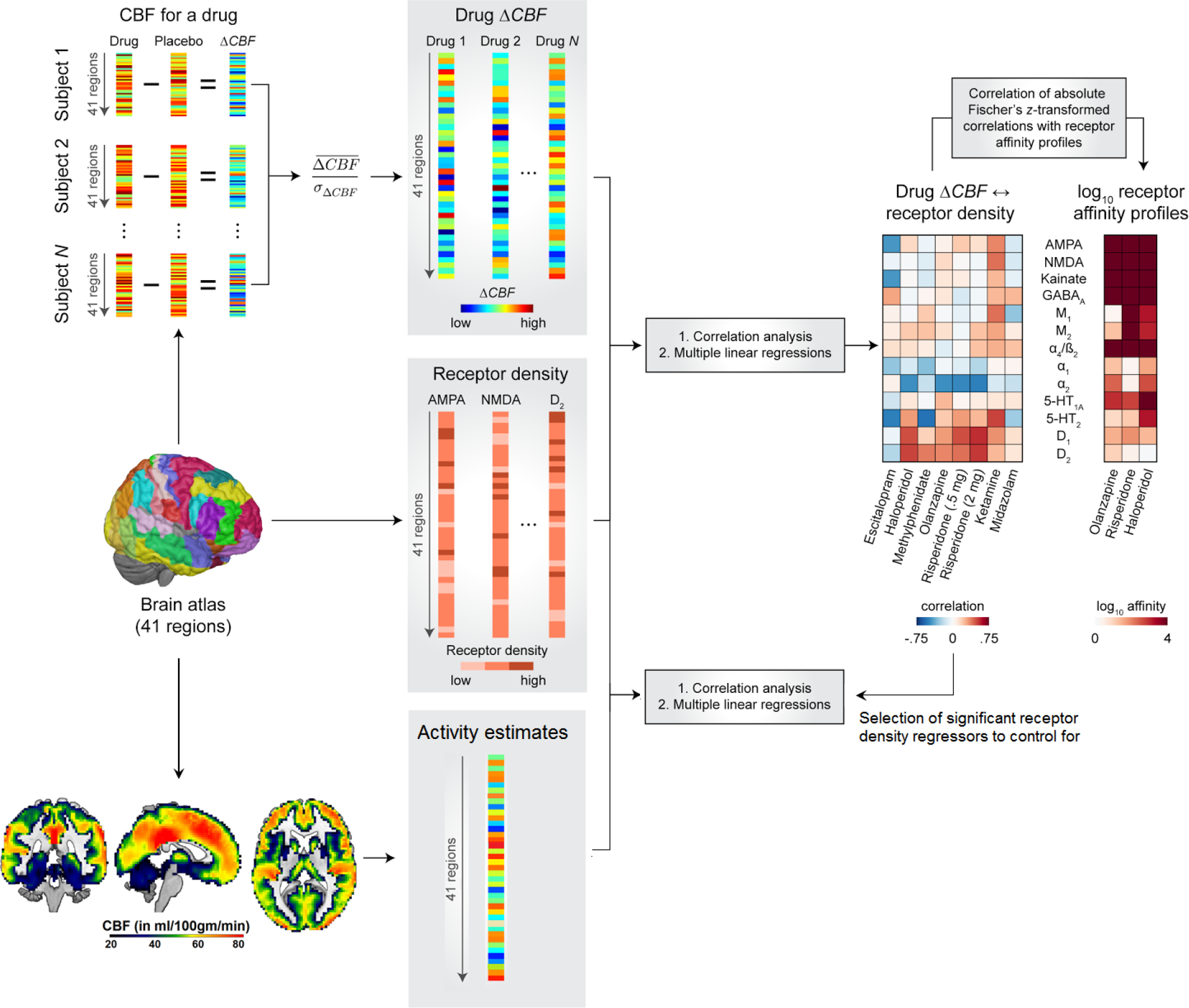
Schematic overview of the proposed mapping of cerebral blood flow (CBF) changes to underlying receptor densities, activity and affinities

**Table 1.** Study data and medication details

|  | Compound |  |  |  |  |  |  |  |
| --- | --- | --- | --- | --- | --- | --- | --- | --- |
|  | Risperidone | Olanzapine | Haloperidol | Methyl-phenidate (Ritalin) | Escitalopram (Lexapro) | Ketamine | Midazolam | Test-retest |
| N subjects | 21 | 21 | 21 | 18 | 18 | 26 | 26 | 29 |
| Study ID | 1 | 2 | 2 | 3 | 3 | 4 | 4 | 5 |
| Demo-graphics (n male, age±SD) | 21, 28±7 | 21, 28±6 | 21, 28±6 | 9, 25±8 | 9, 25±8 | 26, 26±5 | 26, 26±5 | 7, 25±6 |
| ASL acquisitions per session (N) | 2 | 2 | 2 | 1 | 1 | 1 | 1 | 1 |
| Location | KCL | KCL | KCL | UCSD | UCSD | UA | UA | UG |
| Dose (in mg) | 0.5 and 2 | 7.5 | 3 | 30 | 20 | 0.25 mg/kg (bolus), 0.25 mg/kg/hr (infusion) | 0.03 mg/kg (bolus), 0.03 mg/kg/hr (infusion) | - |
| Scanning time (post administration) | 2h | 5h | 5h | 4h | 4h | 0h* | 0h* | - |
| T-max | 1.3h | 4h | 6h | 4.7h | 5h | - | - | - |
| Primary drug indication | SZ, Bipolar | SZ, Bipolar | SZ, Bipolar, Tourette syndrome, Mania | ADHD | Depression, anxiety disorders | Anesthesia, chronic pain | Anesthesia | - |
| Mechanism of action | Direct receptor binding | Direct receptor binding | Direct receptor binding | Reuptake inhibitor | Reuptake inhibitor | Direct receptor binding and reuptake inhibitor | Positive allosteric modulator | - |
| Agonist effects | - | - | - | Dopamine, catecholamines, (serotonin) | Serotonin | Dopamine, norepinephrine, serotonin | GABA | - |
| Antagonist effects | Dopamine, serotonin, catecholamines | Dopamine, serotonin, catecholamines | Dopamine, serotonin, catecholamines | - | - | Glutamate, acetylcholine | - | - |
| Receptors | D1-4, 5-HT 1a and 2a, α1, α2 | D1,D2,D4, 5-HT 1, 2a and 3, D1, α1, α2, muscarinic, H1 | D1-3, 5-HT 2a, α1, σ1 | DAT, NET, SERT | SERT | NMDA, D2, 5-HT 2, AMPA, MAT, nicotinic α4b2, muscarinic | BZD | - |
| Highest affinity to | D2, 5-HT 2a, D3, α2 | 5-HT 2a, D2 | D2, D3 | - | - | NMDA | BZD | - |

ADHD – Attention Deficit Hyperactivity Disorder, AMPA -, ASL – Arterial Spin Labeling, BZD – Benzodiazepine, DAT – dopamine transporter, GABA - Gamma-Amino butyric acid, KCL – King’s College London, MAT – monoamine transporter, NET – norepinephrine transporter, NMDA - N-methyl-D-aspartate receptor, SERT – serotonin transporter, SZ – Schizophrenia, UCSD – University of California San Diego, UA – University of Auckland, UG – University of Groningen, \* intravenous infusion

## Results

### Cerebral blood flow changes correlate with receptor densities

Receptor density maps extracted from a review publication by Palomero-Galagher et al. ^28^ showed a low average co-localization; in addition, compounds with different mechanisms of action also showed low similarity of drug-induced CBF patterns (Figure 2B, Supplementary results, Figure S2 and S3).

**Fig. 2.**
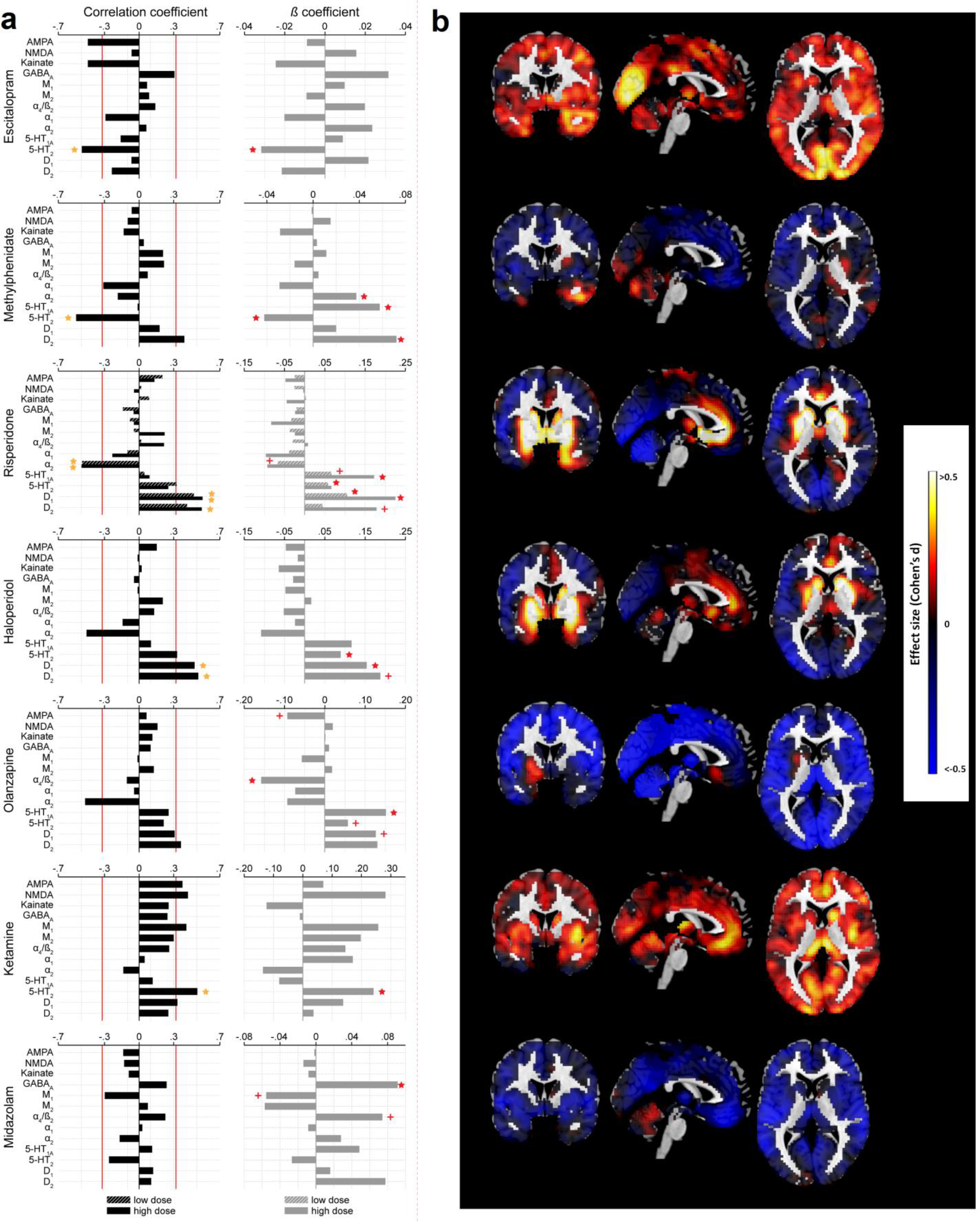
Results of Pearson correlation, multiple linear regression and effect size analyses. a) Results of Pearson correlation (left) and multiple linear regression analyses between receptor densities and CBF changes are displayed as bar plots. For drugs with only one evaluated dose the drug profiles are colored as “high dose”. Red line for Pearson correlation plots indicates significance at an uncorrected two-sided *p*<.05 and yellow star indicates significant Bonferroni corrected findings, For multiple linear regressions a plus indicates a marginally significant (*p*<.1) and red star a significant (*p*<.05) effect of the corresponding regressor. b) Voxel-wise effect size maps (Cohen’s d) are displayed for drug treatments matching the order of drugs displayed in a). For risperidone the outcomes for the high dose are displayed.

We then tested for direct relationships between drug-induced ΔCBF and underlying receptor density maps using correlational analysis. For all compounds beside the GABAergic positive allosteric modulator midazolam we find drug-induced ΔCBF to be consistently correlated with receptor densities corresponding to the known mechanisms of action of the respective drugs (Figure 2A, Table 1). For example, the serotonin agonist escitalopram showed the strongest correlation with serotonergic 5-HT 2 receptor. Consistently, dopamine antagonists risperidone and haloperidol showed significant correlations with D1 and/or D2 receptor densities. Similarly, also olanzapine, methylphenidate and ketamine showed significant correlations (though not all survived Bonferroni correction) with receptor densities corresponding to their known dopaminergic, serotonergic and glutamatergic mechanisms of action. The results of non-parametric analyses using Spearman correlation coefficients were largely similar to the parametric analysis outcomes (Figure S5).

**Fig. 3.**
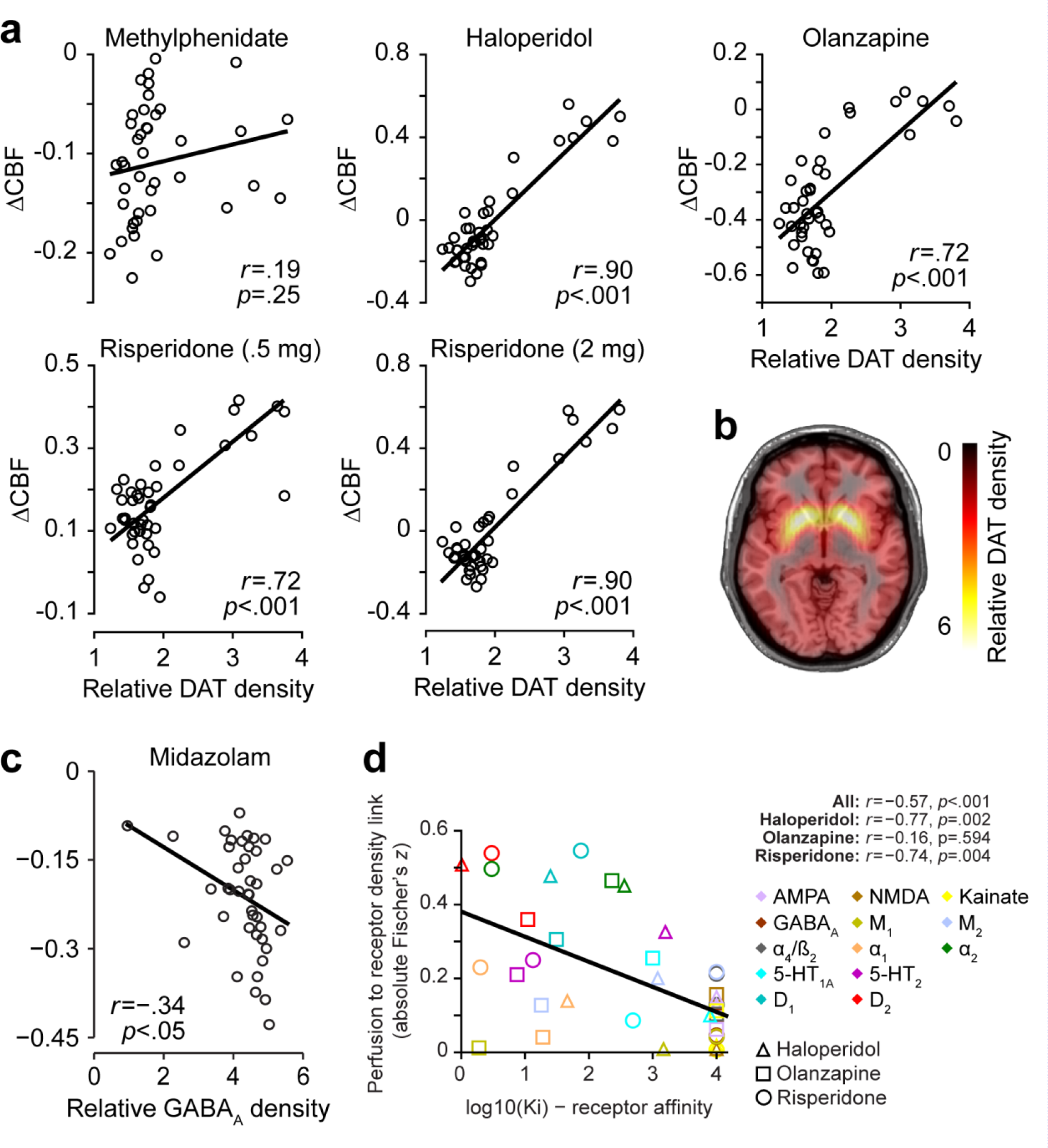
Results of correlational analyses with molecular imaging based receptor density estimates and affinities. a) Correlational plots between regional cerebral blood flow (CBF) changes and respective dopamine transporter (DAT) density profiles are displayed for each drug with dopaminergic mechanism of action. b) DAT density estimates obtained from a healthy volunteer cohort provided by the Parkinson’s Progression Marker Initiative. c) Correlational plot between midazolam induced CBF changes and GABAa density estimates obtained from flumazenil positron emission tomography. d) Correlations of cerebral blood flow (CBF) changes to receptor density profiles with drug affinities. Colors indicate different receptors. Shapes indicate different drugs. Solid line in all plots indicates the linear regression fit.

As the above correlational analyses are based on coarse ex vivo receptor density estimates ^28^, we aimed to evaluate if more fine-grained in vivo density estimates as obtained using molecular receptor imaging of DAT and GABAa are more sensitive for identifying such associations. These correlational analyses revealed very strong and highly significant positive correlations with DAT for the dopamine antagonists haloperidol, olanzapine, and both low and high dose of risperidone but not for the reuptake inhibitor methylphenidate (Figure 3a). Correlation of ΔCBF obtained for the positive allosteric modulator midazolam with flumazenil-based GABAa receptor density estimates revealed a weak but significant association between both (Figure 3c). This correlation remained significant using Spearman correlation coefficient on all data (rho=-.43;p=.005) and after removing the outlier (rho=-.39;p=.013).

### Distinct contribution of different receptor systems to cerebral blood flow changes

Whilst the above correlational analyses provide an estimate of the direct association strength between two measures they do not account for correlations between receptor densities and/or may miss potential weaker associations in presence of a strong effect. Multiple linear regression analyses address these limitations although this is at the cost of a potentially lower sensitivity when regressors are strongly correlated and/or explain similar variance in the dependent variable. To evaluate the distinct contributions of each receptor map to the drug induced ΔCBF and to test for additional associations not discovered through correlational analyses, we therefore computed multiple linear regressions including all receptor densities as regressors in the same model. Significant model fits were observed for all evaluated drugs beside midazolam and the negative control dataset (escitalopram: F(13,27)=2.7;p=.015, haloperidol: F(13,27)=3.2;p=.005, risperidone (low dose): F(13,27)=3.6;p=.002, risperidone (high dose): F(13,27)=5.0;p<.001, olanzapine: F(13,27)=2.8;p=.011, methylphenidate: F(13,27)=3.7;p=.002, ketamine: F(13,27)=2.4;p=.027, midazolam: F(13,27)=1.4;p=.219, negative control: F(13,27)=1.5;p=.175).

To identify which neurotransmitter maps contributed most to these model fits, we then evaluated the significance of each receptor density regressor (Figure 2). These analyses largely replicate the correlational findings and provide evidence of a distinct contribution of the different receptor density maps to the drug-induced ΔCBF. Additionally, they reveal further significant noradrenergic and/or serotonergic contributions for methylphenidate and olanzapine, consistent with their known mechanisms of action. Despite the non-significant overall model fit for midazolam, GABAa was the only significant single receptor density regressor for this compound, in line with midazolam’s mechanism of action.

To assess the robustness of the obtained correlational and multiple regression profiles linking ΔCBF to receptor densities, we evaluated the test-retest reliabilities for compounds where more than one CBF acquisition was available for each subject (risperidone, olanzapine and haloperidol). The association strengths of ΔCBF to receptor density showed excellent test-retest reliability both within and between sessions (intra class correlation coefficients between 0.83 and 0.99; Supplementary results).

### Cerebral blood flow changes correlate with underlying activity

Having established the link between ΔCBF and receptor densities for the respective compounds we then aimed to understand if underlying activity estimates derived from independent non-interventional data are also a predictor of PD effects on CBF. For this we computed correlation coefficient between expected underlying activity and observed ΔCBF for each compound. We observed significant associations between both for midazolam (r=-.48;p=002), methylphenidate (r=-.56;p<.001), haloperidol (r=-.52;p<.001), olanzapine (r=-.62;p<.001), low (r=-.34;p=.028) and high (r=-.57;p<.001) dose of risperidone, a marginally significant effect for escitalopram (r=-.30;p=.055) and no significant effect for ketamine (r=.02;p=.913). When adjusting for the variance explained by receptor densities showing a significant association with respective compounds the effect of underlying activity remained significant for all compounds except escitalopram and ketamine (midazolam: p=.003; methylphenidate: p=.033; haloperidol: p<.001; olanzapine: p<.001; risperidone low dose: p=.050; risperidone high dose: p<.001; escitalopram: p=.471; ketamine: p=.585).

### Strength of CBF to receptor density associations is linked to respective receptor affinities

Whereas the above correlational and multiple linear regression analyses provide evidence of distinct and reliable associations between drug-induced ΔCBF and underlying receptor densities, they remain descriptive with respect to consistency of the observed associations with the respective mechanisms of action of the evaluated compounds. We formally tested this hypothesis by evaluating if the observed association strength profiles between ΔCBF and receptor densities can be explained by drug affinities to the corresponding receptor systems. In a pooled analysis across the three drugs for which the affinity was established using the same methodology we find a highly significant correlation (p<.001) between the obtained profiles and the respective receptor affinities (Figure 3d). Separate tests for each compound confirmed these significant associations for haloperidol and risperidone.

## Discussion

Here we evaluated for seven established compounds acting on dopamine, serotonin, catecholamine, glutamate and GABA the relationship between respective ΔCBF and receptor densities of the underlying neurotransmitter systems. For all compounds, with six out of seven being significant, we find direct and distinct spatial relationships between drug-induced ΔCBF and underlying receptor densities and activity. Moreover, in line with assumptions derived from receptor theory we show that the association strength of these relationships is dependent on the affinity of the drugs to the underlying receptor systems ^27^. Importantly, whilst we test by means of multiple linear regressions for direct linear relationships between ΔCBF and receptor densities, we do not evaluate potential interactions between receptor systems. Due to the often low selectivity the evaluated compounds show high affinity to various receptor systems interactions between neurotransmitter systems are not unlikely. Such interactions may also have resulted in differential CBF responses across regions with different combinations of the targeted receptors and lowered the sensitivity to detect associations between ΔCBF and specific receptor densities. Evaluation of such interactions would yet require larger datasets and more refined receptor density maps.

More specifically, for all evaluated serotonin antagonists we find positive associations between serotonergic system as represented by 5-HT 1a and 2 receptors and drug-induced ΔCBF. These findings suggest that inhibition of this system may be associated with a net increased metabolic demand in the corresponding regions. A more complex picture was observed for serotonin agonists. Whilst ΔCBF induced by both reuptake inhibitors showed a significant negative association with 5-HT 2 receptor density, ketamine showed a significant positive association with this receptor in line with its direct agonist effect on serotonin ^29,30^. This finding suggests interactions between observed ΔCBF and underlying receptor systems, e.g. due to differential effects of orthosteric and allosteric agonists or due to the commonly reported interdependence of dopaminergic, serotonergic and glutamatergic systems ^31–34^

In contrast, all dopamine agonists as well as antagonists evaluated here showed a significant positive association with the underlying D1 and/or D2 receptor densities suggesting a U-shaped association of dopamine levels and metabolic demand. These findings are consistent with previous research reporting increased striatal CBF after administration of both dopamine agonists and antagonists ^35–38^. These findings imply that the hypothesized U-shaped curve relating dopamine to cognitive performance may be paralleled by a U-shaped curve relating dopamine to metabolic demand; for example, cognitive function might improve if dopamine levels are titrated to minimize resting metabolic demands in substrates most strongly associated with those functions ^39,40^. Supportive for these findings with respect to dopamine are also the strikingly strong correlations observed for all three direct dopamine antagonists with in vivo DAT receptor density estimates being a close surrogate of ex vivo counts of dopaminergic neurons ^41^. The strength of these associations with DAT estimates was substantially increased for the high as compared to low dose of risperidone suggesting dose dependency of the observed associations. Supportive for the idea that the established associations between receptor densities and ΔCBF indeed reflect the primary mechanisms of action of the respective compounds are the observed significant correlations of these profiles with respective receptor affinities in the pooled analysis, and also separately for haloperidol and risperidone.

Similarly, the ketamine induced ΔCBF profile was significantly correlated with NMDA (though not surviving Bonferroni correction in the parametric analysis), 5-HT and several other receptor systems including AMPA, for which a ketamine related mechanism of action has only recently been discovered ^30,42^. Though for the GABAergic positive allosteric modulator midazolam the overall model was not significant the finding of GABAa receptor density being the only significant regressor in the multiple linear regression analysis is consistent with its expected mechanism of action. In line with this, *in vivo* estimates of GABAa obtained through flumazenil PET also showed a significant association. Nevertheless, despite reaching significance the association strength with GABAa for midazolam remains weak. This observation is in line with its positive allosteric mechanism of action that would predict its effects to be primarily co-localized with the underlying activity of the system ^43^. Indeed, we find for midazolam and all other compounds with an indirect mechanism of action significant associations between ΔCBF and underlying activity estimates. The strong correlation observed for methylphenidate in that context also explains its weak correlation with DAT to which it directly binds. With a similar argumentation as for midazolam methylphenidate requires activity to take place to exhibit its pharmacological effect. Moreover, in line with our hypothesis we find underlying activity to be also a significant contributor to PD profiles for all compounds with a direct mechanism of action except ketamine. Potential reasons for the lack of such an association for ketamine might be in its wide-spread anesthetic effects that could interfere or alter underlying resting state activity patterns making the applied activity estimates imprecise ^44^.

That said several unexpected findings emerged from our analyses that may relate to limitations of the proposed approach. For example, we do not observe significant associations of escitalopram induced ΔCBF with 5-HT 1a receptor density or between methylphenidate ΔCBF and D1 or DAT receptor density. These negative findings may be false negatives, but may alternatively relate to an important commonality between these compounds: their indirect mechanism of action. Both drugs act as reuptake inhibitors. They therefore require some underlying activity to facilitate its effects. These negative findings may therefore also indicate a relatively low activity of the associated receptor systems at rest. In line with this potential explanation are also the observed significant associations observed for both compounds with the underlying activity estimates. Similarly, despite a high affinity of olanzapine to the M1 receptor we do not observe a significant association between both. This negative finding could suggest a reduced sensitivity of ΔCBF to activity changes related to muscarinic system. However, contrary to this assumption we observe a significant association between M1 and ketamine induced ΔCBF that is also known to have a high affinity to the respective receptor. Among other such discrepant findings may also arise from differences in demographic characteristics between cohorts (i.e. sex ratio) or different scanner types and ASL sequences used for evaluation of different compounds leading to differential sensitivity to detect respective associations.

Differences in observed associations may also arise from the applied drug dosing, frequency, but also weight of the participants. For example, higher dosing may be expected to result in a higher proportion of primary and secondary targets occupied by the drug. This aspect is not unlikely considering that in healthy volunteers dosing was for safety reasons mostly lower as compared to doses applied in clinical routine. Higher variability in weight may increase variability in PK and associated CBF responses. Similarly, some PD effects may evolve on a slower temporal scale. We also observe some significant associations between drug-induced ΔCBF and receptor systems with known low affinity of the respective compounds. In example, a significant association is observed between methylphenidate induced ΔCBF and serotonergic system despite its low affinity to serotonergic receptors ^45^. Such effects are likely to be indirect in nature and may be due to a co-dependence of different networks systems, i.e. through recurrent/long-range interactions with other areas modulated by the compounds ^46,47^. As the applied correlational approach does not differentiate between direct and indirect effects the observed associations may well reflect some previously reported indirect effects of dopaminergic stimulation on the serotonin system ^48,49^. Due to this limitation to correlational relationships with receptor densities, it is also likely that further indirect effects are not detected by the proposed methodology. Lastly, some previous research reported evidence of differential vascular expression of specific dopamine receptors across brain vessels and regions ^20^. This study suggested that such a differential expression may contribute to neurovascular coupling but its translatability to other receptor systems remains unknown. Modulation of such differentially expressed vascular receptors may also have contributed to the relationships between CBF and receptor densities observed in our study. Importantly in that context, the interpretation of our findings with respect to receptor density is limited to a group level setting of drug induced CBF patterns being associated with spatial receptor density profiles associated with the respective compounds. Without further validation the results do not imply any within region relationship across subjects.

In summary, our results provide strong evidence that CBF reflects specific metabolic demands from diverse underlying neurotransmitter systems. We further demonstrate that the proposed approach allows for disentangling of PD effects on CBF to underlying receptor densities and activity, in most cases closely reflecting the mechanisms of action of the respective compounds. These findings support the notion that distinct pharmacological compounds provide unique spatial patterns of CBF changes associated to receptor availability, affinity and function ^1–3,5,24^. With the additionally demonstrated excellent test-retest reliability of obtained CBF to receptor density profiles, these findings further strengthen the value of CBF as a promising tool for drug development and disease evaluation. Our findings and other recent research demonstrate that a combination of pharmacological modulations with neurophysiological read-outs can provide novel insight into specific mechanisms of brain function ^50,51^. Overall, this research shows that the combination of both techniques may provide a unique crossspecies translational approach for studying local and remote neurophysiological and neurometabolic effects associated with modulation of specific neurotransmitter systems. In particular in drug development, the proposed approach may be used to generate data-driven hypothesis about the pharmacodynamic mechanisms of action of novel but also established compounds. Though subject to evaluation the proposed receptor density mapping approach may provide an easy implementable framework for application to other functional neuroimaging measures ^7,16,17,52^ and allows a direct integration of in vivo receptor density estimates as provided by molecular imaging ^53–55^.

## Materials and Methods

### Subject and study details

Data from 5 studies, all in young healthy volunteers, were included. An overview of all studies including medication and dosing details is provided in Table 1. All studies were part of a coordinated effort by F.Hoffmann-La Roche to collect CBF data from drugs with different mechanisms of action following similar fully counter-balanced, placebo controlled, cross-over designs. Adaptations of study designs were performed for each study to account for different administration modes, wash-out times and pharmacokinetic/pharmacodynamic profiles of the evaluated compounds. Study 1 and 2 were conducted using a double-blind, randomized, three-period (each one week apart) cross-over design. Study 1 included three imaging sessions following single dose administration of either low or high dose of risperidone or placebo. Similarly, study 2 included three imaging sessions following single dose administration of olanzapine, haloperidol or placebo. Study 3 was executed in a double-blind, single dose, randomized, four period (each two weeks apart) cross-over design. Evaluated drugs were methylphenidate, escitalopram, a Roche investigational compound (a glutamatergic subtype modulator whose development was terminated, data not reported here) and placebo. Study 4 was a single-blind, randomized, three period cross-over study with imaging performed following intravenous administration of ketamine, midazolam or placebo. Study 5 was a non-treatment test-retest study comprising three MRI visits each including ASL. This study was used as a negative control dataset and to derive activity estimates for the drug studies. Dose selection for all drugs was performed based on the expected receptor occupancy, expected behavioral effects and safety and tolerability considerations. A detailed description of the mechanisms of action for all compounds is provided in Table 1 and Supplement 1. All studies were conducted in accordance with GCP guidelines. Written informed consent was obtained from all study participants. Study 1 and 2 were approved by the London (Brent) Human Research Ethics committee (REC reference: 13/LO/1183). Study 3 was approved by the Institutional Review Board at the University of California, San Diego (OMB No. 0910-0014). Study 4 was approved by the Health and Disability Ethics Committees of the Ministry of Health, Wellington (Ethics ref.: 15/CEN/254). Study 5 was approved by the local Ethics Committee in Assen, Netherlands (Stichting Beoordeling Ethiek Biomedisch Onderzoek).

### ASL data

Image acquisition onset for each study was based on the expected PK maximum for the corresponding compounds (2 hours post dose for Study 1, 5 hours post dose for Study 2, 4 hours post dose for Study 3). ASL data with varying sequences) were acquired for all studies and all visits among other study specific imaging modalities. For Study 1 and 2 two runs of ASL were acquired at each session. For these studies, run 2 data of each session were used for primary analyses to reduce potential arousal effects at the beginning of the acquisition. Details on ASL acquisition and preprocessing of resulting cerebral blood flow data are provided in Supplement 1 (Supplementary methods, Table S1 and Figure S6). In brief, CBF computation for studies 1, 2, 4 and 5 was based a pseudo-continuous ASL acquisition whilst a FAIR QUIPSS II ASL sequence was used for study 3. For all studies, CBF was computed in standard physiological units (ml blood/100mg tissue/min) based on sequence specific recommendations ^11,56^. All pseudo-continuous sequences included acquisition of a proton density image to enable appropriate CBF quantification. Due to lack of identical ASL sequences for different manufacturers and scanner types, the best available scanner-specific ASL protocol at the time of study conduct was used for each study.

### Receptor density maps

Receptor density maps for 41 regions (Brodmann areas and subcortical nuclei) were extracted for the following 13 receptor types from the publication by Palomero-Gallagher et al.^28^: for glutamate (AMPA, NMDA and Kainate), for GABA (GABAa), for acetylcholine and muscarine (M1 and M2), for nicotine (nicotinic α4/β2), for catecholamines (α1 and α2), for serotonin (5-HT 1A and 5-HT 2), for dopamine (D1 and D2) (Table S2). A 3 point coarse scale provided by the authors was applied for all receptor systems, e.g. 1=low, 2=intermediate, 3=high, For some regions intermediate levels of receptor densities between those 3 levels were reported. Those were coded as 1.5 or 2.5. Brodmann regions have been shown to provide distinct functional information as measured through resting state MRI if at all rather under-parcellating such data ^57^. Additionally, as for each receptor map for about 5% of regions the density was reported as unknown we aimed to reduce data loss for the multiple linear regression analyses requiring a full data matrix. For this an interpolated version of the receptor density table was created replacing the missing values by the mode of other densities for the corresponding receptor. A detailed description of receptor density map extraction is provided in Supplement 1.

As these coarse receptor density maps were obtained from a review publication of ex vivo studies, we aimed to additionally evaluate if more fine grained in vivo density estimates provided by molecular receptor imaging further improve the observed associations. For this we extracted dopamine transporter (DAT) and GABAa density estimates as measured through DAT-SPECT and flumazenil PET for the 41 regions reported above. DAT-SPECT data were obtained from a publicly available control cohort of healthy volunteers (Parkinson’s Progression Marker Initiative) (Figure 3B). GABAa density estimates were obtained using flumazenil PET data of 6 healthy volunteers acquired at the Imperial College London. Details on these cohorts, pre-processing and DAT and GABAa density estimation are provided in Supplement 1.

### Pharmacodynamic CBF profiles

A Brodmann area map as included in the MRIcron tool ^58^ was normalized into the Montreal Neurological Institute space using the Statistical Parametric Mapping (SPM12) normalize function ^59^. For putamen, caudate and fusiform gyrus regions the corresponding automated anatomical labeling atlas regions were used. Mean CBF values were extracted for each subject for drug and placebo images from the 41 regions corresponding to Brodmann and subcortical areas covered by the receptor density maps described above. A delta drug minus placebo (change versus placebo) was then computed for each subject and region. A ΔCBF profile for each drug was calculated as an effect size per region for the whole group by computing the average regional drug-induced change across all subjects divided by the standard deviation of the change in the respective region across all subject (Figure 2B and Supplementary Figure 4). Major differences in receptor densities and pharmacological effects across individuals have been reported in earlier animal studies that are also likely to apply to human experiments ^60,61^. Effect size normalizes the mean signal in each region by the variability of the signal. Therewith, it takes this variability into account whilst providing an index of the drug effect. It was therefore chosen to minimize the potential impact of such between subject and region variability in signal to noise but also due to site, scanner and sequence differences across different studies. For Study 5, test retest data of each subject for visit 1 and 2 were randomly assigned to either drug or placebo condition. All group-level ΔCBF imaging data alongside with receptor density maps and activity estimates are provided in Supplementary material.

### *Mapping* ΔCBF *to receptor densities*

For interpretation of subsequent correlations between receptor density and ΔCBF maps we first aimed to understand if and how the obtained receptor maps provide similar or differential information regarding spatial receptor density distribution. Pearson correlations and coefficients of determination were computed between each pair of receptor density maps to estimate their association strength. Validity of the parametric statistics described below (parametric and normality assumptions for Pearson correlations and multiple linear regression models) was established using permutation statistics and Shapiro-Wilk tests for normality (s. Supplementary methods and results for description and outcomes of those analyses, Figure S1). Additionally, to further ensure that the choice of parametric tests did not bias the results we repeated all analyses using Spearman correlation coefficients (Figure S5).

We next aimed to understand if and how receptor density maps are linked to observed drug-induced ΔCBF profiles (1) separately for each receptor type and (2) whilst controlling for the variance explained by other receptor density maps. For these we performed two types of analyses: (1) simple pair-wise Pearson correlations between each receptor density map and each ΔCBF map and (2) multiple linear regression models using each ΔCBF map as a dependent variable and including all 13 receptor maps as regressors in a single model. To ensure that the outcomes of analysis (1) are not biased by coarse receptor density scale, outliers or distribution assumptions we recomputed all correlations using Spearman correlation coefficients (Figure S5). All analyses were implemented using default parameters in IBM SPSS Statistics (Version 23.0, IBM Corp., Armonk, NY). For analysis (1) we report if correlations survive strict Bonferroni correction (accounting for the number of tests performed per compound) and additionally as exploratory findings all correlations surviving an uncorrected p<.05. For analysis (2) to reduce data loss due to missing values the interpolated receptor density maps described in Supplement 1 were used. To ensure that the interpolation did not bias the results, this analysis was repeated using the initial receptor density maps excluding regions with missing values (s. Supplementary results). In analysis (2) we test for significance of each overall model and each single receptor density regressor to predict ΔCBF profiles (p<.05) whilst controlling for the effects of all other regressors included in the model.

We then tested if continuous in vivo receptor density estimates further improve the associations between receptor densities and ΔCBF. For this we computed for all compounds with known dopaminergic mechanism correlation coefficients with DAT density estimates obtained through DAT-SPECT. Similarly, we computed for midazolam being the only GABAergic compound its correlation with flumazenil-based GABAa receptor density estimates. To ensure that the putative outlier region showing a very low GABAa expression (Figure 3c) did not bias the results we further repeated the correlation analysis with GABAa using non-parametric Spearman correlation coefficient with all data and after the removing the outlier.

Lastly, as several ASL acquisitions were available for some of the compounds, we used those data to estimate the reliability of the obtained ΔCBF to receptor density and affinity mappings: (1) within session data of ASL run 1 and 2 for haloperidol, olanzapine and risperidone and (2) between session data for low and high dose data of risperidone. For both, within session and between session data we then used intra-class correlation coefficients (ICC(C,k)) to assess reliability of established receptor density to ΔCBF profiles (s. Supplement 1 for detailed outcomes of those analyses).

### Testing for associations with underlying activity

To evaluate if drug-induced ΔCBF is also linked to underlying activity we used the non-drug study 5 data from all visits to compute a quantitative average CBF map (Figure 1). Mean CBF values across all subjects extracted from this map for the 41 regions introduced above were used as independent expected activity estimates for the respective regions for all drug data. Similarly to the above associations with receptor density we followed a two-step procedure to test if drug-induced ΔCBF is associated with expected activity profile. In a first analysis, we tested for associations between activity profile and ΔCBF using Pearson correlations. We then tested using multiple linear regressions if these relationships still hold when controlling for the contributions of significant receptor density regressors identified above.

### Testing for associations with receptor affinity

To further understand if the observed relationships between receptor densities and ΔCBF are also associated with the respective receptor affinities (Ki) we first extracted the affinities for compounds with direct receptor binding mechanism of action established using the same methodology (risperidone, olanzapine and haloperidol) from Bymaster et al. (Table S3) ^62^. For receptors with no detectable drug binding (Ki>10000 nM) the value 10000 was used. If affinities to several receptor subtypes were reported the average affinity of these subtypes was used. For further analyses receptor binding affinities for all drugs were converted to log values (e.g. log(10000)=4) ^63^. Further, we hypothesized that if ΔCBF are indeed related to the underlying receptor systems, stronger (positive or negative) correlations should be observed between both for receptors with higher drug affinities. To test this assumption we used the absolute Fisher’s z transformed Pearson correlation coefficients obtained in analysis (1) for each of the 3 drugs and each receptor density map (13 values). These association strength profiles between receptor densities and ΔCBF were then correlated with the respective receptor affinities for each drug (pooled and separately for each compound).

## Acknowledgments

DAT-SPECT data used in the preparation of this article were obtained from the Parkinson’s Progression Markers Initiative (PPMI) database (www.ppmi-info.org/data). For up-to-date information on the study, visit www.ppmi-info.org. PPMI – a public-private partnership – is funded by the Michael J. Fox Foundation for Parkinson’s Research and funding partners, including Abbvie, Avid Radiopharmaceuticals, Biogen Idec, Briston-Myers Squibb, Covance, GE Healthcare, Genentech, GlaxoSmithKline, Lilly, Lundbeck, Merck, Meso Scale Discovery, Pfizer, Piramal, Roche and UCB.

## Author contributions statement

JD designed the study. JD and FS wrote the manuscript. JD, SH, CC, PH, AF, RM, JM, ARL, DJN, EM, CR, LB, DU, SS, TL, MAM, FOZ, SCW, GB, MP, GDH, SM, JH, AB and FS contributed to design and execution of individual studies. All authors reviewed and commented on the manuscript.

## Data availability statement

All group level drug and receptor density maps used in the study are provided as supplementary material.

